# solPredict: Antibody apparent solubility prediction from sequence by transfer learning

**DOI:** 10.1101/2021.12.07.471655

**Authors:** Jiangyan Feng, Min Jiang, James Shih, Qing Chai

**Author notes:** Jiangyan Feng; Qing chai.

## Abstract

There is growing interest in developing therapeutic mAbs for the route of subcutaneous administration for several reasons, including patient convenience and compliance. This requires identifying mAbs with superior solubility that are amenable for high-concentration formulation development. However, early selection of developable antibodies with optimal high-concentration attributes remains challenging. Since experimental screening is often material and labor intensive, there is significant interest in developing robust *in silico* tools capable of screening thousands of molecules based on sequence information alone. In this paper, we present a strategy applying protein language modeling, named solPredict, to predict the apparent solubility of mAbs in histidine (pH 6.0) buffer condition. solPredict inputs embeddings extracted from pretrained protein language model from single sequences into a shallow neutral network. A dataset of 220 diverse, in-house mAbs, with extrapolated protein solubility data obtained from PEG-induced precipitation method, were used for model training and hyperparameter tuning through five-fold cross validation. An independent test set of 40 mAbs were used for model evaluation. solPredict achieves high correlation with experimental data (Spearman correlation coefficient = 0.86, Pearson correlation coefficient = 0.84, R^2^ = 0.69, and RMSE = 4.40). The output from solPredict directly corresponds to experimental solubility measurements (PEG %) and enables quantitative interpretation of results. This approach eliminates the need of 3D structure modeling of mAbs, descriptor computation, and expert-crafted input features. The minimal computational expense of solPredict enables rapid, large-scale, and high-throughput screening of mAbs during early antibody discovery.

## Introduction

Therapeutic monoclonal antibodies (mAbs) represent the fastest growing class of therapeutics on the market, with around 100 antibody drugs approved to treat a wide spectrum of human diseases ^1^, including cancer ^2,3^, inflammatory and auto-immune diseases ^4^. Subcutaneous injection has emerged to be the preferred delivery route of mAbs drug products especially in the treatment of chronic diseases, because they can be self-administered at home and therefore enhances patient adherence and compliance ^5^. Given limited injection volume (<2 mL) and high dose requirement (∼500 mg), mAbs must be soluble enough to achieve high-concentration formulations (>100 mg/mL) ^6^. Furthermore, mAbs must remain soluble at high concentrations during the manufacturing process which can cause protein precipitation. Therefore, superior solubility is vital for developing liquid formulation of therapeutic mAbs ^7–9^.

A practical hurdle is that poor solubility behavior often manifest at higher mAb concentrations (>50 mg/mL) ^10^. Early experimental screening is often challenged by the large number of antibody candidates and the limited preparation quality available (i.e. minute amounts, low concentrations, and low purity) ^10,11^. *In silico* solubility prediction appears to be a convenient alternative owing to its capability of rapid high-throughput screening without material requirement ^12–14^. Current computational approaches rely on molecular descriptors extracted either from protein sequence (sequence-based predictors ^13,14^) or from structures (structure-based predictors ^12,15^). Sequence-based predictors often neglect tertiary structure information, which distinguishes poorly soluble residues driving protein folding from the ones that are exposed to the solvent and may elicit aggregation ^7^. Structure-based tools can be used only when the structure or a high-quality model are available. This limits the throughput and application to large number of early stage mAb candidates. Furthermore, some of the computational methods only output a binary classification (e.g. soluble/insoluble) ^13,16,17^ instead of a numerical value.

The lack of quantitative solubility dataset of large, diverse mAbs at pharmaceutically relevant formulation further hinders the generalizability of computational predictors. Previous developability related work has been performed with non-mAbs proteins ^13^, limited mAb datasets ^12,14^, closely related mAbs with varying mutations ^12,14^, or mAbs belonging to the same subclass ^18^. Furthermore, mAb solubility is highly dependent on formulation condition ^10^. Histidine and pH 6.0 (H6) buffer system has emerged as a common buffer/pH system for mAb-based products, because at pH 6.0 chemical degradation of proteins is minimized which makes liquid formulations feasible ^6^. To the best of our knowledge, there are no computational tools that can quantitatively predict the solubility at H6 condition for different subclasses of mAbs using sequence information alone.

A powerful approach is transfer learning which distills the knowledge learned from protein language models trained on very large unlabeled protein sequences and builds a downstream model supervised with limited data ^19^. Previous studies have shown that using the representations from pretrained protein language models as input features to fine-tune a supervised model can improve a wide variety of protein-relevant tasks, including secondary structure prediction, contact prediction, and remote homology detection ^20,21^. However, such methods have not been fully explored to predict attributes that are essential to antibody developability.

Here, we propose an end-to-end sequence-to-function model, solPredict, to predict the quantitative solubility of mAbs at H6 using only antibody sequences. To develop a general predictor of antibody solubility, we constructed a large and diverse set of 260 in-house mAbs, consisting of 112 IgG1, 140 IgG4, and 8 IgG2. The quantitative solubility was measured at histidine pH 6.0 buffer condition using PEG-induced precipitation method due to the advantages of high-throughput screening and minimal material requirement ^10^. To overcome the limitation of expert-crafted descriptors, and the necessity to obtain high-quality 3D structures, we represented antibody sequences as embeddings, fixed-length vectors extracted from a pretrained protein language model (ESM1b) ^20^. We show that pretrained protein embeddings are informative for mAbs property prediction. Supervised learning using simple machine learning models and small labeled dataset suffice to enrich the signals and learn the sequence-to-solubility relationship. We also find that mAb solubility behavior differs among different IgG subclasses. solPredict can predict IgG1 and IgG4 reliably, but more IgG2 data is needed to be generalizable to IgG2. In this work, we provide a systematic framework for the study of antibody developability when limited data is available, which can be used for high-throughput screening of mAbs during early antibody discovery.

## Results

### Dataset construction

We sought to construct a large, diverse dataset with quantitative measures of solubility behavior. We used polyethylene glycol 3350 (PEG 3350) with concentration ranging from 4 to 36% to assess the relative solubility of 260 mAbs (see *Materials and methods* for details). We chose the PEG-induced approach due to its capability of robust, high-throughput screening with minimal material consumption ^10^. Out of the 260 mAbs, 112 are IgG1, 140 are IgG4, and 8 are IgG2 molecules. Measurements were made in a 0.4 M L-histidine (pH 6.0) buffer condition, which has emerged as a popular choice for mAb-based products ^6^. To investigate the intrinsic experimental noise, we examined two control mAbs with known solubility behaviors (mAb239: IgG1, good solubility; mAb240: IgG4, poor solubility) (Table 1). The two control mAbs were measured under histidine buffer pH 6.0 (H6) condition for 47 times. The summary statistics are shown in Table 1 and the distribution of solubility measurements is reported in Figure 1a. The measurement of the well behaved control (mAb239) is 35.81 ± 0.72 PEG % while the measurement of the poorly behaved control (mAb240) is 10.63 ± 1.46 PEG %, demonstrating the robustness of the PEG-induced approach to differentiate mAbs by their solubility.

**Table 1.** Summary of solubility behavior of two control mAbs at H6.

| mAb | Subclass | Solubility<br>behavior | H6 (PEG %) |  |
| --- | --- | --- | --- | --- |
|  |  |  | mean | standard<br>deviation |
| mAb239<br>(control 1) | IgG1 | Good solubility | 35.81 | 0.72 |
| mAb240<br>(control 2) | IgG4 | Poor solubility | 10.63 | 1.46 |

**Figure 1.**
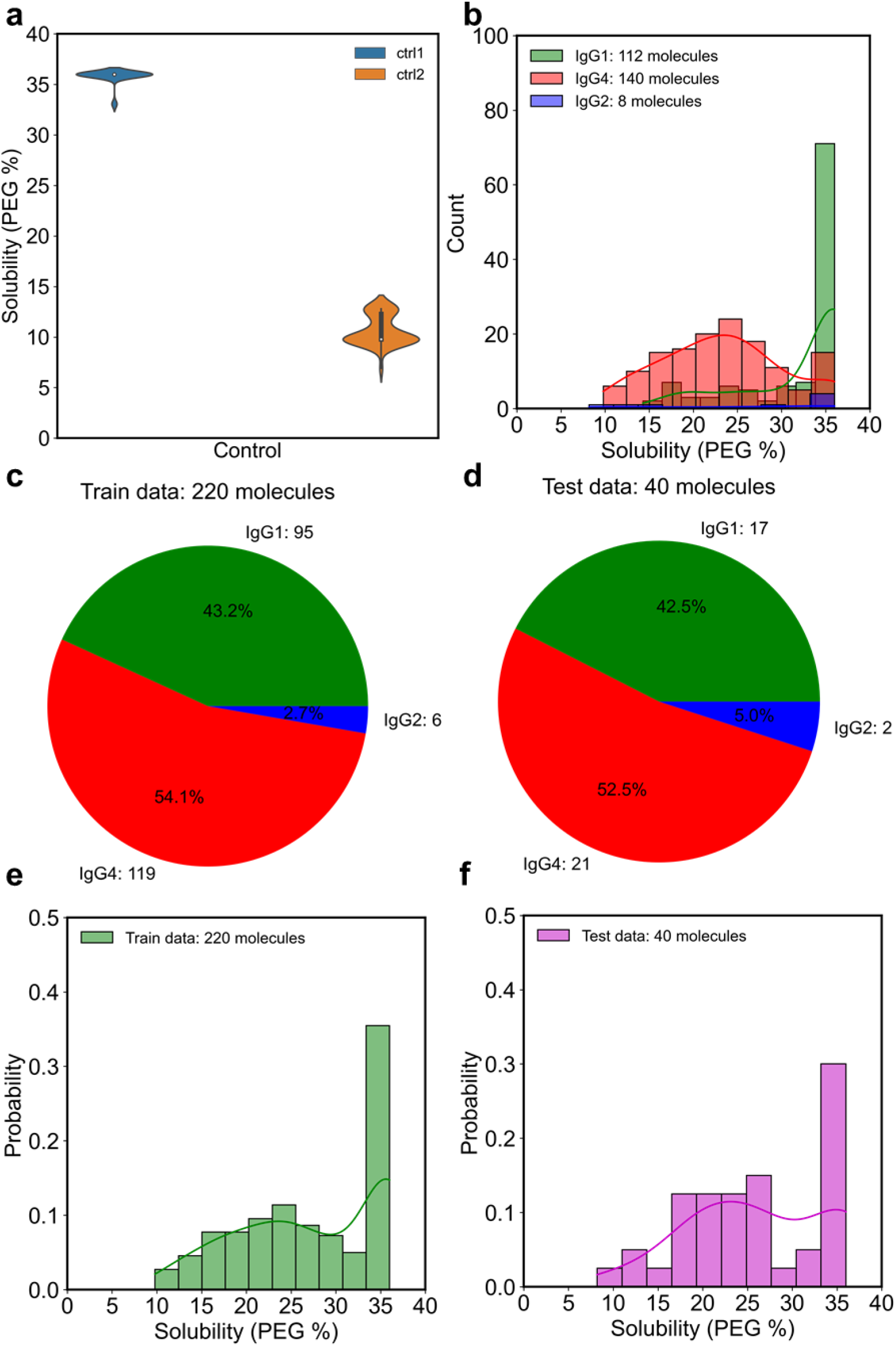
mAb solubility dataset. (a) The solubility distribution of two controls at histidine buffer pH 6.0 (H6). For each control, the solubility measurements were repeated for 47 times. (b) The solubility distribution of different IgG subclasses. The IgG subclass composition for (c) training dataset (n=220) and (d) test dataset (n=40). The solubility distribution for (e) training dataset (n=220) and (f) test dataset (n=40).

Next, we explored the contribution of antibody subclass to solubility behavior (Figure 1b). Previous developability studies focused on the variable domains due to the sequence and structural homology between IgG subclasses ^22,23^. However, our results suggest IgG1 and IgG4 exhibit divergent solubility behavior. A broad range of solubility was observed in IgG4 molecules, whereas IgG1 mAbs tend to be highly soluble under H6 condition. Similarly, previous study reports that IgG1 and IgG4 mAbs show totally different behavior in terms of viscosity, opalescence, and thermostability ^6,24,25^. Therefore, a dataset consisting of different subclasses is critical to the development of computational predictors that can be generalizable to diverse mAbs. To ensure the similar subclass composition of train/test dataset, we randomly split the dataset into train and test group (85/15) for each subclass (Figure 1c, d, e, f). The train dataset contains 220 mAbs, consisting of 95 IgG1, 6 IgG2, and 119 IgG4 (Figure 1c). The test dataset contains 40 mAbs, composed of 17 IgG1, 2 IgG2, and 21 IgG4 (Figure 1d). The final solubility distribution is similar between train and test set (Figure 1e, f).

### Pretrained protein language model embeddings enable the use of small, labeled dataset and simple architecture for mAb solubility prediction

While the current dataset of 260 mAbs is already larger than previous mAbs developability studies ^6,22,23,25–29^, it is still smaller than most machine learning projects in which millions of data points were used for training ^30^. To overcome the data scarcity, we propose an end-to-end machine learning framework for solubility prediction with transfer learning from pretrained protein language model (ESM-1b) ^20^ (Figure 2). ESM-1b model was trained on 86 billion amino acids across 250 million unlabeled protein sequences in an unsupervised manner. The learned representations contain rich information about biological properties, ranging from biochemical properties of amino acids to the secondary and tertiary structure. Our proposed framework is mainly empowered by the informative protein representation from ESM-1b, which captures the general sequence semantics in the protein universe. At first, we extracted embeddings for both heavy and light chains from the last layer of ESM-1b. Second, we used the concatenated embeddings (2560-dimensional feature vector, 1280 for each chain) as input and trained various downstream regression models for solubility prediction.

**Figure 2.**
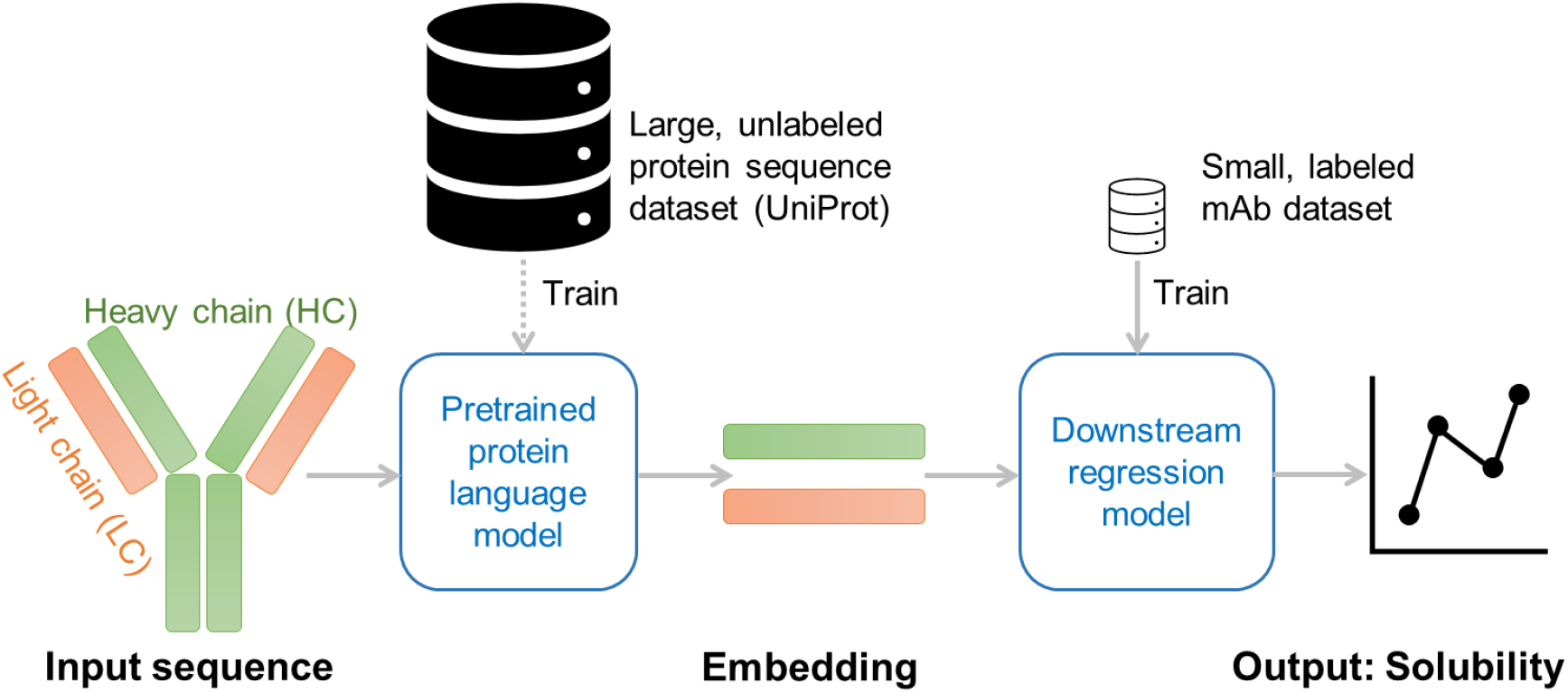
The solPredict architecture. First, full IgG sequences were converted into fixed size embeddings (1280 for each chain) through protein language model pretrained using large, unlabeled protein sequence database. Next, the heavy chain and light chain embeddings were concatenated into 2560-dimensional feature vectors and used as the input of a downstream regression model that predicts mAb solubility. Quantitative solubility data measured by PEG-induced precipitation method were used to supervise the training of the regression model.

We compared traditional machine learning models (support vector machine regressor (SVM), random forest regressor (RF)) to neutral network models (multilayer perceptron (MLP) with 1 fully connected hidden layer (MLP1Layer), MLP with 2 fully connected hidden layers (MLP2Layer)). For SVM and RF, the input embeddings were first compressed into 23-dimensional vectors using principal component analysis (PCA) to explain > 90% variance. For MLP1Layer and MLP2Layer, the input embeddings were directly used as the first layer of the network. Five-fold cross validation of the training dataset of 220 mAbs was used for hyperparameter tuning for all models. Once the optimal hyperparameter set was determined, the final models for SVM and RF were trained on the whole training set using these hyperparameters. For MLP1Layer and MLP2Layer model, Spearman correlation coefficient on validation set was used to select the best checkpoint during training. The average Spearman correlation coefficient was used to select the optimal combination of hyperparameter and the resulting five models. Instead of refitting with the whole training set, the mean of five models was used for prediction. The hyperparameter search range and the selected combination of hyperparameter is shown in Table S1 and Figure S1.

The test dataset of 40 mAbs were used to compare these four models (Figure 3, Table S2, Figure S2). We reported a variety of performance metrics for systematic comparisons: Spearman correlation coefficient, Pearson correlation coefficient, R^2^, and root mean square error (RMSE). Despite the simplicity of architecture, all four models achieved high correlation with experimental solubility data. This suggests that pretrained embeddings are effectively informative that simple machine learning models supervised with a small annotated experimental dataset suffice to predict antibody solubility with high performance. For instance, the SVM model achieves comparable performance with the more complex MLP1Layer model (Spearman correlation coefficient: 0.80 versus 0.81, Pearson correlation coefficient: 0.79 versus 0.81, R^2^: 0.61 versus 0.64, RMSE: 4.89 versus 4.72) (Table S2). Although RF model outperforms the others on the training dataset (Figure S2), its performance is the worst on the test set (Figure 3, Table S2), indicating overfitting issue. Adding 1 more hidden layer further boosts the performance, with Spearman correlation coefficient increases from 0.81 to 0.86, Pearson correlation coefficient increases from 0.81 to 0.84, R^2^ increases from 0.64 to 0.69, and RMSE reduces from 4.72 to 4.40. This may be because MLP2Layer is better at learning the complex sequence-to-solubility relationship. No additional hidden layer was tested due to the limited dataset. As more quantitative solubility data become available, more complex neutral networks may further improve the performance. As MLP2Layer model performs the best based on all four evaluation metrics, it was selected as the downstream regression model for solPredict. Taken together, we show that protein language model pretrained on general protein sequences can provide a powerful signal for antibody related downstream task.

**Figure 3.**
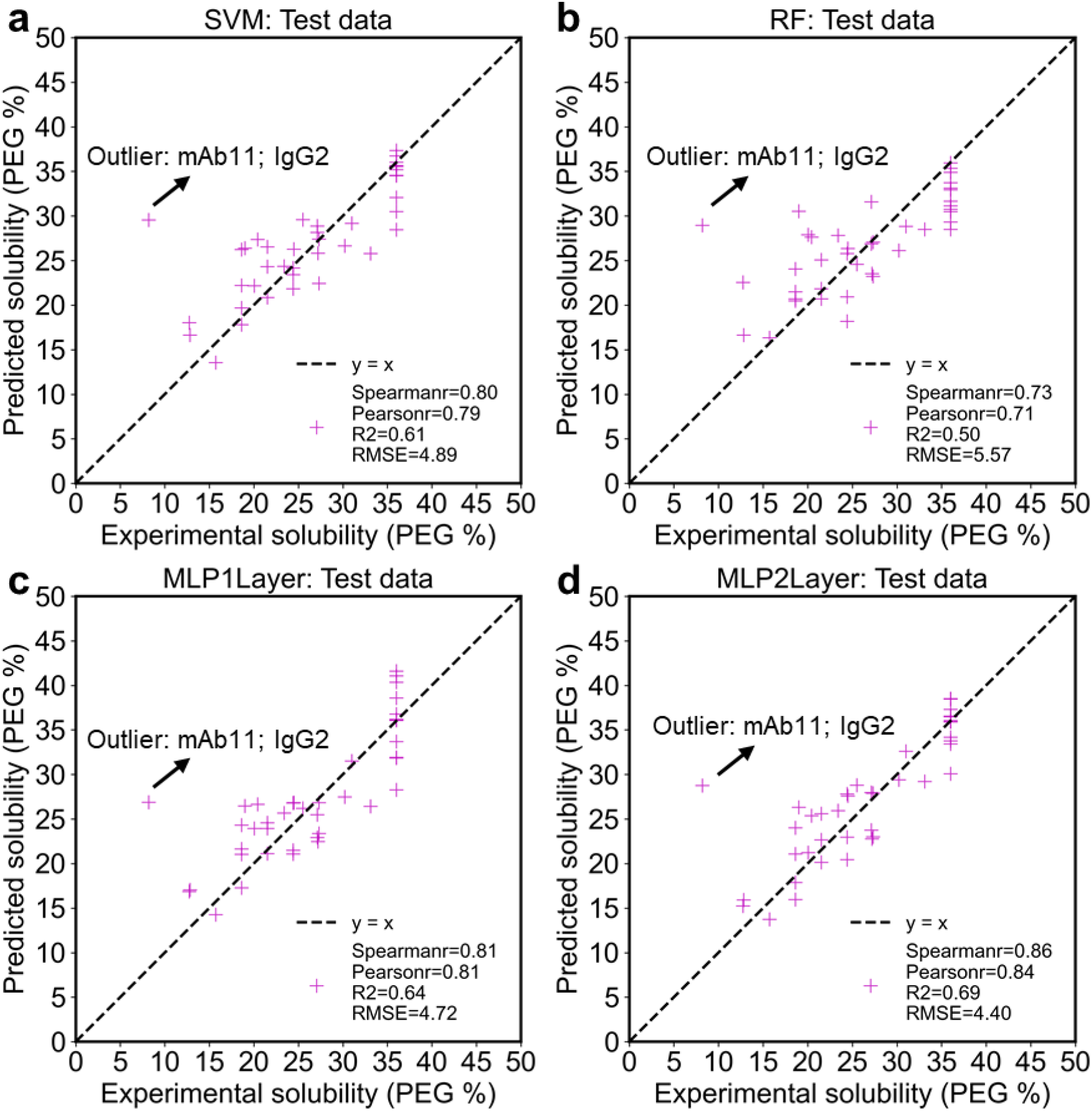
Performance of different regression models on test dataset (n=40). The correlation between the predicted (y-axis) and experimentally measured solubility (x-axis) on the test dataset for (a) SVM, (b) RF, (c) MLP1Layer, and (d) MLP2Layer models. The dashed black line refers to the perfect correlation: y=x. The statistics of four evaluation metrics are shown in legend. The outlier (mAb11, IgG2) is annotated for each model.

### solPredict can predict both IgG1 and IgG4 accurately

Next, we investigated how performance varies among different isotypes (Figure 4). Our analysis shows that solPredict can predict both IgG1 (n=17) and IgG4 (n=21) with high performance (Figure 4a, b, Table S3). The performance on the IgG1 test set is the best with Spearman correlation coefficient = 0.84, Pearson correlation coefficient = 0.93, R^2^ = 0.87, and RMSE = 2.36. The performance on the IgG4 test set is slightly worse than IgG1 with Spearman correlation coefficient = 0.69, Pearson correlation coefficient = 0.77, R^2^ = 0.58, and RMSE = 3.48. This is due to the different solubility behavior between IgG1 and IgG4 molecules. Most IgG1s exhibit high solubility (>30 PEG %), while IgG4s exhibit a bell curve distribution, with the bulk showing medium solubility (∼25 PEG %) (Figure 1b, Figure 4a, b). The relationship between sequence and solubility is therefore simpler to learn for IgG1 compared with IgG4 molecules.

**Figure 4.**
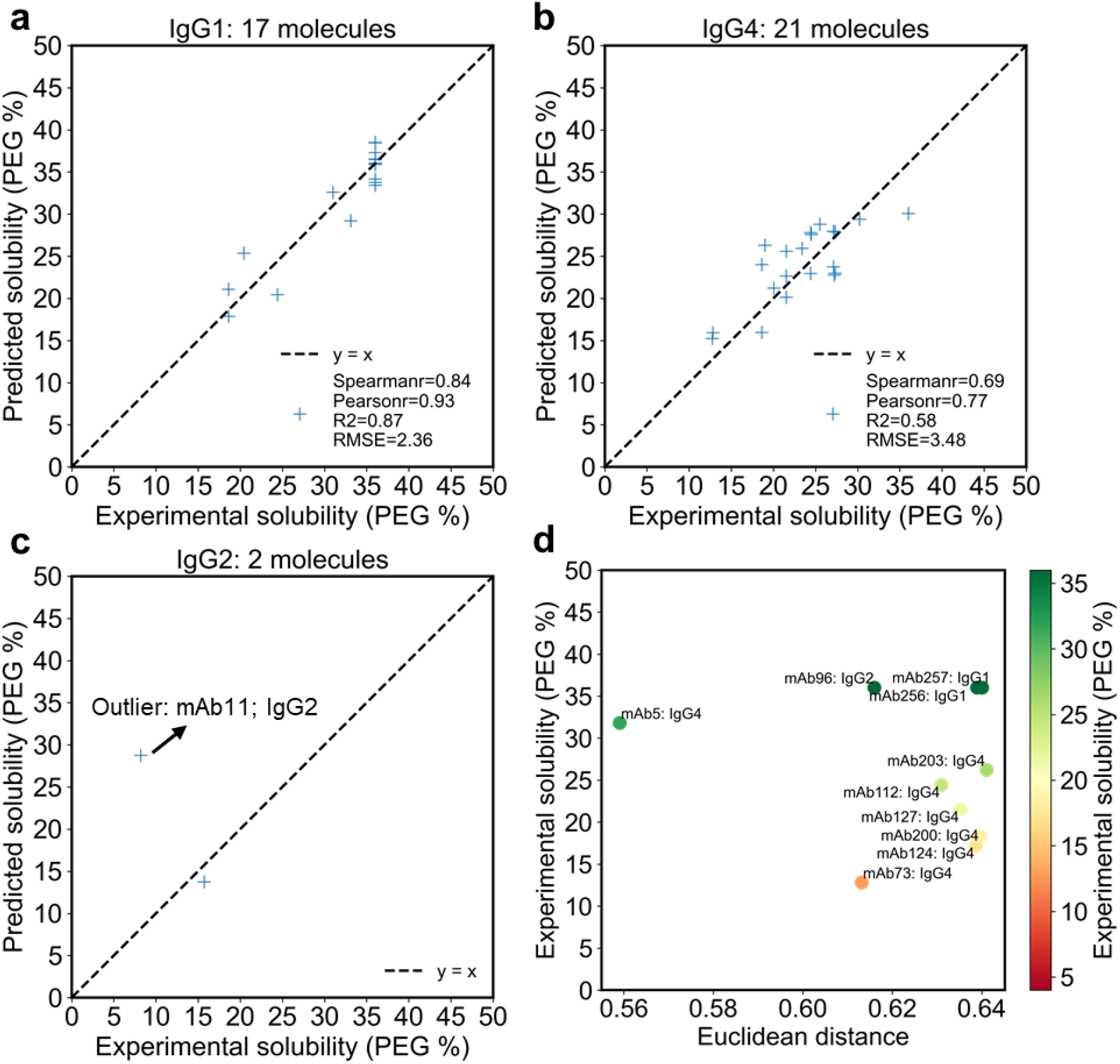
Performance of solPredict (MLP2Layer as downstream regressor) on different IgG subclasses. The correlation between the predicted (y-axis) and experimentally measured solubility (x-axis) for (a) IgG1 test dataset (n=17), (b) IgG4 test dataset (n=21), and (c) IgG2 test dataset (n=2). The dashed black line represents the perfect correlation: y=x. The values of four evaluation metrics are shown in legend for IgG1 and IgG4. Due to the limited sample size of IgG2 (n=2), no evaluation metric was computed. The outlier (mAb11, IgG2) is highlighted in (c). (d) Visualization of 10 training molecules closest to mAb11. The x-axis represents the Euclidean distance of embeddings between mAb11 and the other mAbs. The y-axis represents the experimentally determined solubility. Each data point is colored based on the experimentally determined solubility and annotated by the mAb index and subclass.

For IgG2 (n=2), there is an outlier, mAb11, whose solubility was overestimated by all four models (Figure 4c; Figure 3a, b, c, d). The experimental solubility for this IgG2 molecule is 8.20 PEG %, whereas all four models consistently predicted it around 30 PEG %. We sought to understand why mAb11 can’t be reliably predicted. We first compared the embeddings of mAb11 against all 220 training molecules and selected the top 10 mAbs that are closest to mAb11 based on the Euclidean distance (Figure 4d, Table 2). 7 out of the 10 closest neighbors are IgG4 and the closest neighbor is mAb5, an IgG4 molecule with high solubility (31.80 PEG %). Only 1 out of the 10 closest neighbors is IgG2, which also shows high solubility (36.00 PEG %). It appears that solPredict predicts mAb11 to be much more soluble than the experimental measurement because it reasons with the mapping relationship learned from IgG4 molecules. It suggests that the sequence-to-solubility mapping relationship for IgG2 is different from IgG1 and IgG4, and solPredict couldn’t learn the sequence-to-solubility mapping for IgG2 due to the limited IgG2 molecules (n=6) in the training set. To sum up, solPredict can serve as an accurate predictor of IgG1 and IgG4 mAb solubility, while more IgG2 data is needed to ensure the generalizability to IgG2 prediction.

**Table 2.** Top 10 mAbs in the train dataset closest to mAb11 ranked by the Euclidean distance.

| mAb | Euclidean distance | Subclass | H6 (PEG %) |
| --- | --- | --- | --- |
| mAb11 | 0 | IgG2 | 8.20 |
| **mAb5** | **0.56** | **IgG4** | **31.80** |
| mAb73 | 0.61 | IgG4 | 12.80 |
| **mAb96** | **0.62** | **IgG2** | **36.00** |
| mAb112 | 0.63 | IgG4 | 24.40 |
| mAb127 | 0.64 | IgG4 | 21.50 |
| mAb124 | 0.64 | IgG4 | 17.15 |
| mAb256 | 0.64 | IgG1 | 36.00 |
| mAb200 | 0.64 | IgG4 | 18.30 |
| mAb257 | 0.64 | IgG1 | 36.00 |
| mAb203 | 0.64 | IgG4 | 26.20 |

### Pretrained protein embeddings are informative for antibody solubility behavior

To interrogate how solPredict learns to predict mAb solubility, we projected the embeddings into two dimensions with t-distributed stochastic neighbor embedding (t-SNE) ^31^. We performed the analysis both for the raw embeddings from ESM1b (Figure 5a) and for the final hidden layer of the MLP2Layer network after training (Figure 5b). The key hyperparameter (perplexity) was optimized for t-SNE construction (see *Materials and methods* for details). All t-SNE representations in Figure 5 were created using 5000 iterations, PCA as initialization, and Euclidean distance as similarity metric. The perplexity was set as 10 for the raw representation and 30 for the last hidden layer representation. The t-SNE representations were colored according to their experimental solubility at H6. Although never trained, the raw embeddings appeared to capture some information about mAb solubility, with diffuse organization of small communities sharing similar solubility behavior (Figure 5a). After training, the network learned the sequence-to-solubility relationship and showed clear structure of representation space with the diagonal corresponding to the variation in solubility (Figure 5b). The outlier mAb11 was incorrectly placed in the middle of high solubility mAbs, further verifying our previous hypothesis that the relationship of IgG2 was not well learned yet. Together, it shows that pretrained protein embeddings contain general properties about protein sequences and further supervised learning with a small dataset suffice to reorganize the embeddings for specific task.

**Figure 5.**
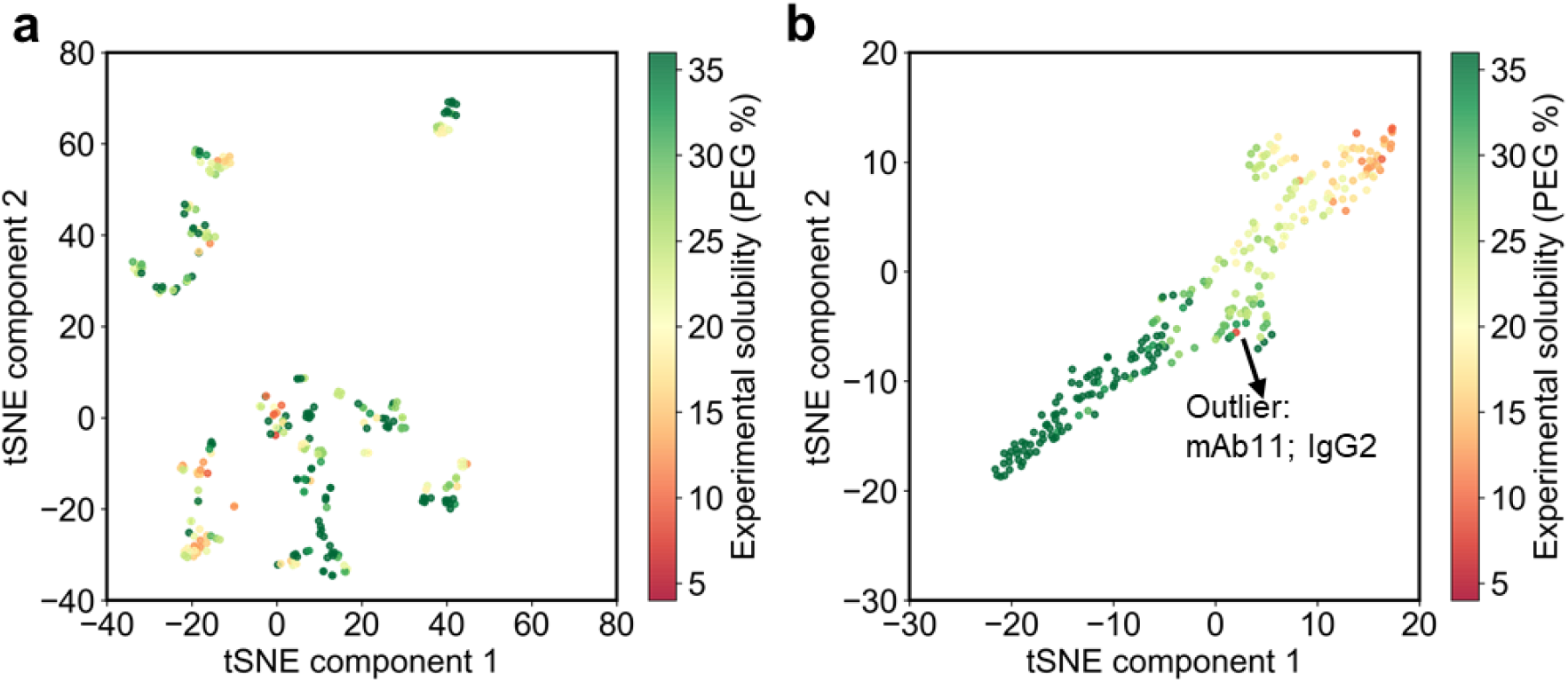
Visualization of embeddings along two dimensions using t-SNE. The representation of all 260 mAbs using (a) raw embeddings and (b) last hidden layer of MLP2Layer model after training. The outlier (mAb11, IgG2) is highlighted in (b). Each point represents a mAb and each mAb is colored by the experimentally measured solubility.

## Discussion

We propose that incorporating high-capacity protein language model pretrained on 100s of millions of sequences stored in protein databases (e.g. UniProt ^32^) will be the key to alleviate both the data scarcity and feature engineering challenges for antibody developability prediction. As a proof-of-concept, we applied the pretrained protein language model to predict mAb solubility through transfer learning. We represented full IgG sequences using embeddings extracted from the pretrained protein language model. Using the embeddings as input, we trained four simple machine learning models (SVM, RF, MLP1Layer, and MLP2Layer) with a diverse set of 220 mAbs. Methods were compared using an independent test set of 40 mAbs. We find that all four models show high correlation with experimental measurements despite the simplicity of models. MLP2Layer model performs the best with Spearman correlation coefficient = 0.86, Pearson correlation coefficient = 0.84, R^2^ = 0.69, and RMSE = 4.40. Furthermore, MLP2Layer model can predict both IgG1 and IgG4 mAbs with high performance despite their distinct solubility behavior. Our results suggest that pretrained protein embedding are powerful representations for sequence-to-property mapping for mAbs. Further supervised learning with small, labeled dataset can enhance the signal for specific task, solubility prediction in this case. We anticipate that transfer learning and massive protein language models can also be used to predict other developability properties, such as viscosity, aggregation propensity, and stability.

solPredict can reliably predict antibody apparent solubility from sequence information alone, which makes it suitable for screening the solubility of a large library of mAbs during early antibody discovery. In general, PEG <= 19 % is used as guideline for flagging mAbs with poor solubility in the screening assay. With 19 PEG % as threshold, solPredict achieves a successful classification rate of ∼90% on the test dataset (Figure S3a). The errors mainly come from false positives. 4 out of 40 mAbs were misclassified as high soluble (1 IgG1, 2 IgG4, and 1 IgG2 molecules) while none of them was misclassified as low soluble (Figure S3a, b, c, d). It suggests that solPredict will be less likely to eliminate otherwise highly soluble mAb candidates, which is more costly and consequential than false positive error.

solPredict can also be integrated into existing protein engineering workflows to guide the discovery of beneficial variants enhancing solubility. For example, diverse variants (single/double mutants or subtype switch) can be rapidly predicted using solPredict. Promising mutations can be combined for the next rounds of predictions. After a few rounds of optimization, the best variants that maximize the solubility can be prioritized for experimental testing, in which the screened data can be used to further improve the model, hereby forming an iterative loop of computational prediction and experimental validation to discover enhanced variants.

In summary, our work confirms the value of pretrained protein language models on antibody properties and demonstrates a new framework for rapid and high-throughput antibody developability prediction using only sequence information.

## Materials and methods

### Antibodies

A total of 260 in-house mAbs, consisting of 112 IgG1, 140 IgG4, and 8 IgG2 subclass were used in this study. They were all produced internally at Eli Lilly & Co using either HEK293 or Chinese hamster ovary cells expression system. mAbs were purified using a standard antibody purification procedure (protein A capture followed by polishing steps). All reagents and excipients were commercially available from Hampton Research, EM Chemicals, JT Baker, Sigma-Aldrich, and Mallinckrodt, and were of high purity (> 98%).

### PEG-induced precipitation for solubility measurement

The apparent solubilities of mAbs were measured using PEG-induced precipitation assays, as described in a previous publication ^10^. Stock solutions of 0.4 M L-histidine, pH 6.0 were used for H6 buffer matrices. Solutions with Polyethylene glycol 3350 (PEG 3350) levels varying from 4 to 36 % were prepared and were mixed overnight at 1200 RPM at 25°C. All mAbs were buffer-exchanged and diluted with water to a target final concentration of 1 mg/mL. After mixing mAbs and stock solutions on each assay plate, all assay plates were sealed with foil and incubated at 25°C on the bench for 24 hours. Finally, the plates were read at 280 nm (with background subtraction at 320 nm) using a Tecan Infinite M1000 Pro UV/Vis Spectrophotometer. The absorbance data was de-convoluted and plotted using Excel. The onset of precipitation was determined visually based on the point of abrupt decrease in absorbance caused by the loss of protein. The nearest PEG 3350 concentration (%) corresponding to the onset of precipitation was used as the estimation of solubility. Therefore, the experimental measurement of solubility varies from 4 to 36 PEG %. The higher the PEG %, the better is the solubility.

### Protein embeddings computation

ESM1b model ^20^ was used as the pretrained protein language model to extract the embeddings for mAbs. ESM1b model was trained on 86 billion amino acids across 250 million protein sequences using unsupervised learning and improved a range of applications such as prediction of mutational effect, secondary structure, and long-range contacts. These representations therefore contain both sequence level and structural level signals. For each input sequence, ESM1b generates a 1280-dimensional vector representation for each amino acid. In this work, the average of all amino acid features was used as a final feature vector for each sequence (1280-dimensional vector per sequence). The full sequence of heavy chain and light chain for each IgG was used to generate 1280-dimensional vector per chain. The heavy chain and light chain embedding were then concatenated into a 2560-dimensional vector to represent each mAb.

### Training details

For all four models, 220 mAbs were used for training and hyperparameter tuning through five-fold cross validation. For support vector machine and random forest regressor, we used PCA to reduce the dimensionality of the raw embeddings from 2560 to 23 to explain 90% of variance. Hyperparameters were optimized using a grid search and five-fold cross validation (Table S1). Once the hyperparameters were optimized, the whole training dataset of 220 mAbs was used to refit the support vector machine and random forest regressor. Scikit-learn ^33^ was used to implement support vector machine and random forest models.

The neutral network models (MLP1Layer and MLP2Layer) were trained in a mini-batch mode using Pytorch ^34^. MLP1Layer refers to multilayer perceptron with 1 fully connected hidden layer. MLP2Layer refers to multilayer perceptron with 2 fully connected hidden layers. ReLU was used as the activation function. The training process takes 500 epochs on the training dataset of 220 mAbs using the Adam optimizer and Xavier_uniform initialization. The hyperparameters (batch size, learning rate, and hidden layer dimensions) were tuned using five-fold cross validation (Table S1). The training objective is to minimize the mean squared error between predicted and experimental solubility values. To alleviate overfitting, the Spearman correlation coefficient on the validation set (the held-out fold) was used to select the best checkpoint during training. The average of Spearman correlation coefficient on validation sets was used to select the best combination of hyperparameters and the final set of models. The selected hyperparameters for MLP1Layer model are hidden layer size = 256, batch size = 32, and learning rate = 0.01. The selected hyperparameters for MLP2Layer model are hidden layer 1 size = 64, hidden layer 2 size = 32, batch size = 8, and learning rate = 0.001.

### Performance evaluations

The test set of 40 mAbs were never used for model training or hyperparameter tuning. The performance on test set were evaluated using Spearman correlation coefficient, Pearson correlation coefficient, R^2^, and RMSE. Spearman and Pearson correlation coefficient were implemented using SciPy python package ^35^. Both correlation coefficients vary between -1 and +1 with 0 implying no correlation. The difference is that Pearson correlation assumes that data is normally distributed whereas Spearman correlation is a nonparametric measure without assuming that datasets are normally distributed. Considering the solubility data distribution is skewed in the dataset, Spearman correlation was used as the main evaluation metric. Scikit-learn ^33^ was used to compute R^2^ (sklearn.metrics.r2_score) and RMSE (square root of mean squared error: sklearn.metrics.mean_squared_error). R^2^ (coefficient of determination) measures how well the regression predictions approximate the real data points with 1 indicating the perfect fit. R^2^ can be negative if the model performs worse than a constant model (R^2^ = 0). RMSE is a measure of the difference between predicted and actual values, with 0 indicating the perfect fit between predicted and actual data points. The lower the RMSD, the better the regression model.

### t-SNE construction

To project the high-dimensional embeddings on to two-dimensional space, we used the t-distributed stochastic neighbor embedding (t-SNE) algorithm implemented in Scikit-learn ^33^. Perplexity parameter was varied from 5 to 100 with a step size of 5. Similarity metric was based on Euclidean distance. PCA was set as initialization and 5000 iterations were performed. In the end, the perplexity was selected as 10 for the raw representation and 30 for the last hidden layer representation.

## Acknowledgments

We thank Peter Hillier and Yuhao Lin for valuable software support. We thank Guilherme Rocha, William Smith, and Ahmadreza Ghanbarpour Ghouchani for helpful discussion. We thank the antibody engineering group and automation group at Lilly Biotechnology Center for characterizing, producing, and purifying the antibodies.

## Abbreviations

H6: histidine buffer pH 6.0 (H6)
mAb: monoclonal antibody
MLP: multilayer perceptron
PCA: principal component analysis
RF: random forest
RMSE: root mean square error
SVM: support vector machine
t-SNE: t-distributed stochastic neighbor embedding
PEG: polyethylene glycol

