## Supplemental material for "solPredict: Antibody apparent solubility prediction from sequence by transfer learning"

Table S1 Hyperparameter search for downstream regressors.

| Regressor | Hyperparameters | Selected hyperparameter |
|---|---|---|
| SVM | C: [0.1, 1.0, 10.0]<br>kernel: [‘linear’, ‘poly’, ‘rbf’, ‘sigmoid’]<br>degree: [3]<br>gamma: [‘scale’] | C: [10.0]<br>kernel: [‘rbf’]<br>degree: [3]<br>gamma: [‘scale’] |
| RF | # of estimators: [20]<br>criterion: [‘mse’, ‘mae’]<br>max features: [‘sqrt’, ‘log2’]<br>min samples split: [5, 10]<br>min samples leaf: [1, 4] | # of estimators: [20]<br>criterion: [‘mse’]<br>max features: [‘log2’]<br>min samples split: [5]<br>min samples leaf: [1] |
| MLP1Layer | # of epochs: [500]<br>hidden layer size: [64, 128, 256, 512]<br>batch size: [4, 8, 16, 32]<br>learning rate: [0.001, 0.01] | # of epochs: [500]<br>hidden layer size: [256]<br>batch size: [32]<br>learning rate: [0.01] |
| MLP2Layer | # of epochs: [500]<br>hidden layer 1 size: [64, 128, 256]<br>hidden layer 2 size: [32, 64, 128]<br>batch size: [8, 16, 32]<br>learning rate: [0.001, 0.01] | # of epochs: [500]<br>hidden layer 1 size: [64]<br>hidden layer 2 size: [32]<br>batch size: [8]<br>learning rate: [0.001] |

Table S2 Results comparison of different regression models on test dataset.

| Regressor | Spearman | | Pearson | | $R^2$ | RMSE |
|---|---|---|---|---|---|---|
| | correlation | p value | correlation | p value | | |
| SVM | 0.80 | 7.13e-10 | 0.79 | 1.27e-09 | 0.61 | 4.89 |
| RF | 0.73 | 6.88e-08 | 0.71 | 2.73e-07 | 0.50 | 5.57 |
| MLP1Layer | 0.81 | 2.37e-10 | 0.81 | 2.19e-10 | 0.64 | 4.72 |
| **MLP2Layer** | **0.86** | 1.67e-12 | **0.84** | 1.72e-11 | **0.69** | **4.40** |

Table S3 Performance of MLP2Layer on IgG1 and IgG4 subclasses.

| Subclass | Sample size | Spearman | | Pearson | | $R^2$ | RMSE |
|---|---|---|---|---|---|---|---|
| | | correlation | p value | correlation | p value | | |
| IgG1 | 17 | 0.84 | 2.35e-05 | 0.93 | 3.93e-08 | 0.87 | 2.36 |
| IgG4 | 21 | 0.69 | 5.18e-04 | 0.77 | 4.38e-05 | 0.58 | 3.48 |

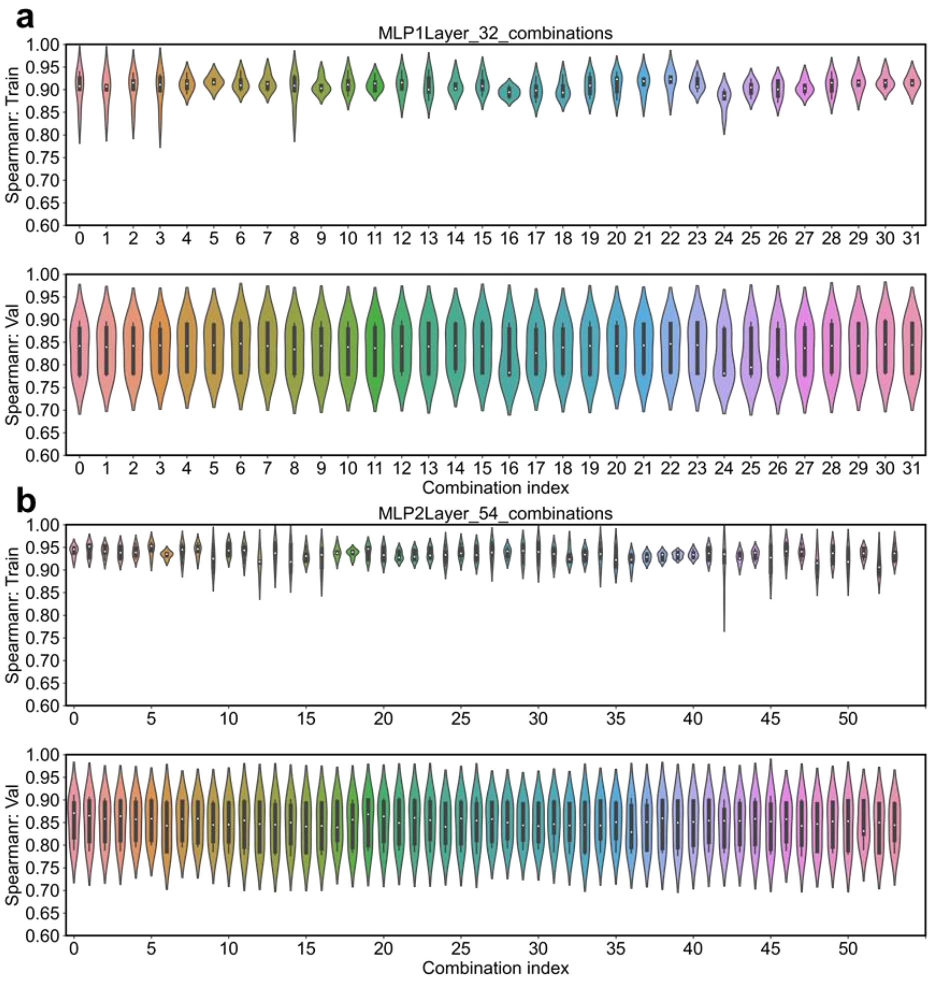


Figure S1 Hyperparameters search results for MLP1Layer and MLP2Layer. Five-fold cross validation on the training set of 220 mAbs was used to tune hyperparameters. For each set of hyperparameters, 5 models were constructed. Each model was trained using 4 folds of training dataset and evaluated using the remaining fold (validation dataset). The average Spearman correlation on the validation dataset was used to select the best combination of hyperparameters. The Spearman correlation coefficient distribution on the training set and validation set is shown for (a) MLP1Layer and (b) MLP2Layer. In the end, combination #30 was selected for MLP1Layer, with the best average validation Spearman correlation as 0.84. The hyperparameters are hidden layer size = 256, batch size = 32, and learning rate = 0.01. Combination #0 was selected for MLP2Layer, with the best average validation Spearman correlation as 0.86. The hyperparameters are hidden layer 1 size = 64, hidden layer 2 size = 32, batch size = 8, and learning rate = 0.001.

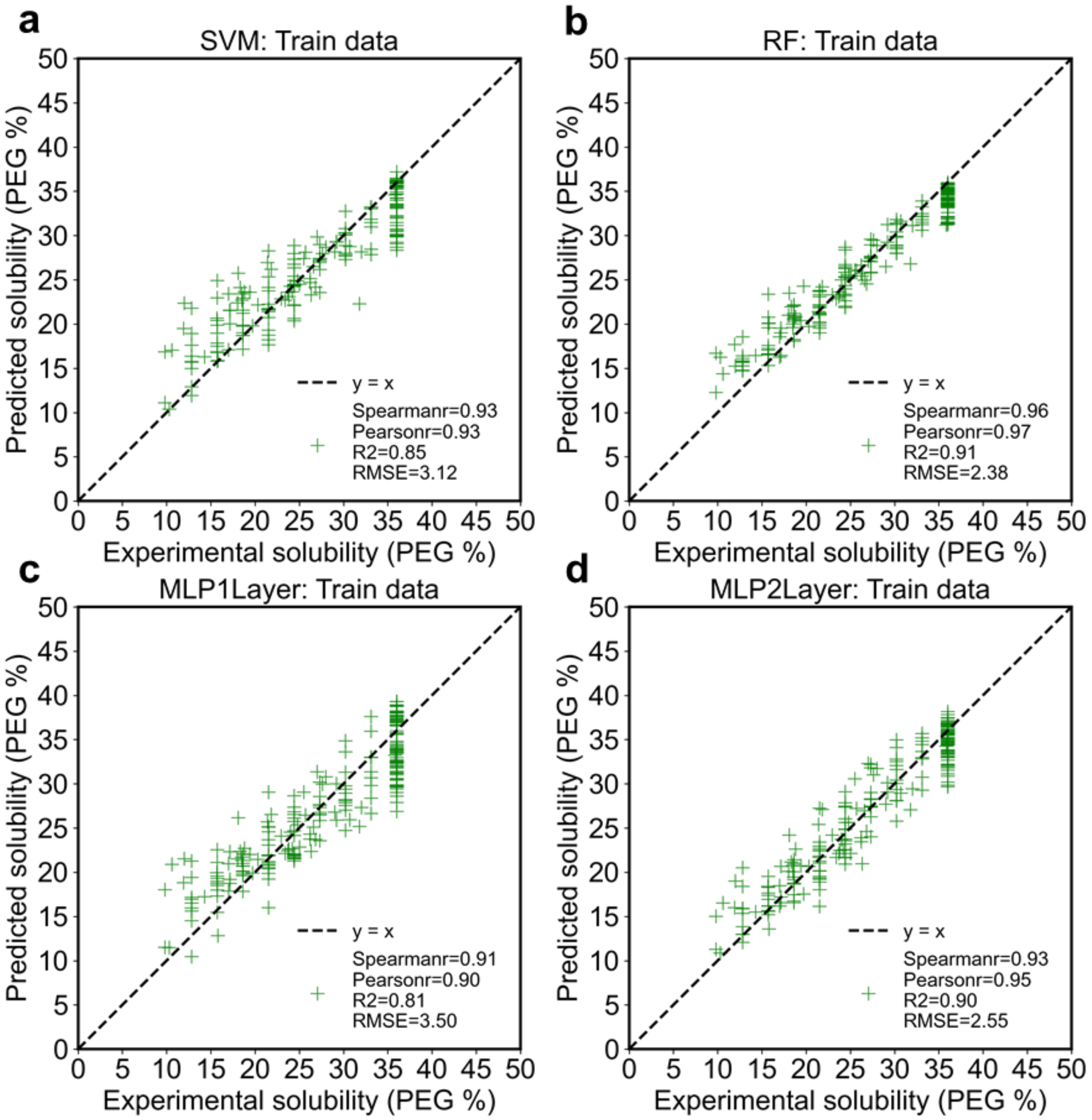


Figure S2 Performance of different regression models on train dataset (n=220). The correlation between the predicted (y-axis) and experimentally measured solubility (x-axis) on the train dataset for (a) SVM, (b) RF, (c) MLP1Layer, and (d) MLP2Layer models. The dashed black line refers to the perfect correlation: y=x. The performance on four evaluation metrics is shown in legend. It is important to note that SVM and RF models were refit using the whole train dataset (n=220) after hyperparameter selection, whereas MLP1Layer and MLP2Layer models were not refit.

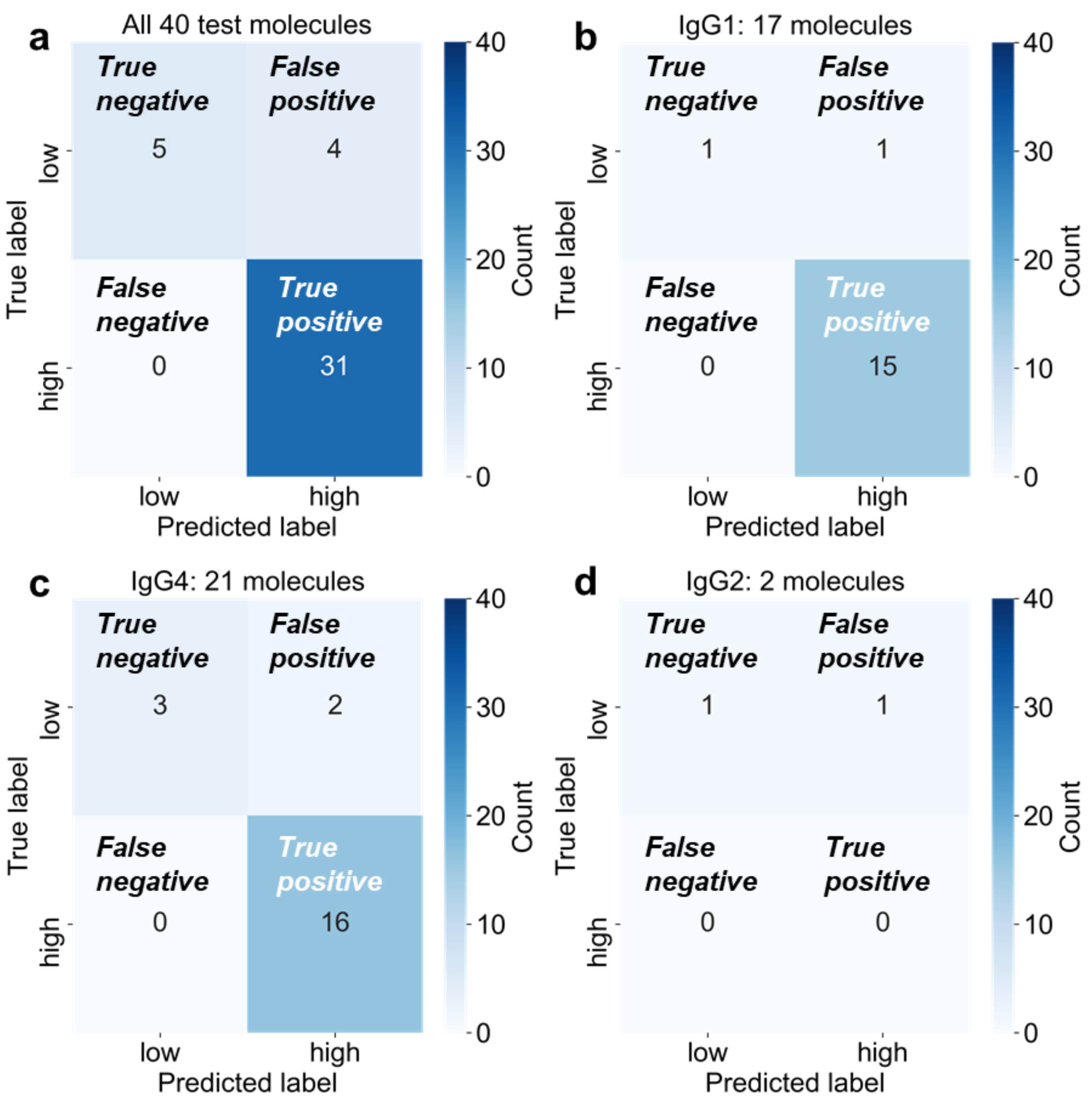


Figure S3 Performance of solPredict on binary classification. The quantitative output from solPredict can be adapted for binary classification with specified threshold. 19 PEG % was used as the threshold to differentiate high (> 19 PEG %) and low (<= 19 PEG %) solubility. The confusion matrix is shown for (a) the whole test dataset (n=40), (b) IgG1 test dataset (n=17), (c) IgG4 test dataset (n=21), and (d) IgG2 test dataset (n=2). True negative, true positive, false negative, and false positive are annotated in specific regions and were colored by counts. 5 of 40 test molecules were correctly predicted as low soluble. 31 of 40 test molecules were correctly predicted as high soluble, giving an overall accuracy of 90%.
